# Degenerated virulence and irregular development of *Fusarium oxysporum* f. sp. *niveum* induced by successive subculture

**DOI:** 10.1101/2020.10.29.356675

**Authors:** Tao-Ho Chang, Ying-Hong Lin, Yu-Ling Wan, Kan-Shu Chen, Jenn-Wen Huang, Pi-Fang Linda Chang

## Abstract

Successive cultivation of fungus has been reported to cause the sectorization, which leads to degeneration of developmental phenotype, and virulence. *Fusarium oxysporum* f. sp. *niveum* (Fon), the causal agent of watermelon Fusarium wilt, demonstrated that successive cultivation formed the degenerated sectors. In the present research, we demonstrated that subculture with aged mycelium increased the incidence of degenerations. To further investigate the differences between the Fon wild type (sporodochial type, ST) and variants (MT: mycelium type and PT: pionnotal type), developmental phenotypes and pathogenicity to watermelon were examined. Results have shown that degeneration variants (PT2, PT3, PT11 and MT6) were different from ST with mycelium growth, conidia production and chlamydospore formation. Virulence of degenerated variants on susceptible watermelon Grand Baby (GB) cultivar was determined after inoculated with Fon variants and Fon ST. In root dipping methods, all Fon variants showed slightly increased disease severity than ST. Conversely, all Fon variants showed a significant decrease in disease progression compared with ST through infested soil inoculation. The contrary results of two inoculation methods suggest that the changes of successive cultural degeneration may lead to the loss of pathogen virulence-related factors of the early stage of Fon infection process. Therefore, Cell wall-degrading enzymes (CWDEs; cellulase, pectinase, and xylanase) activities of different variants were analysed. All Fon degenerated variants demonstrated significantly decreased of CWDEs activities compared with ST. Additionally, transcripts level of 9 virulence-related genes (*fmk1*, *fgb1*, *pacC*, *xlnR*, *pl1*, *rho1*, *gas1*, *wc1 and fow1*) were assessed in normal state. The degenerated variants demonstrated a significantly low level of tested virulence-related genes transcripts except for *fmk1*, *xlnR* and *fow1*. In summary, the degeneration of Fon is triggered with successive subculture through aged mycelium. The degeneration showed significant impacts on virulence to watermelon, which caused by the reduction of CWDEs activities and declining expression of a set of virulence-related genes.

## 1. Introduction

*Fusarium oxysporum* (Fo), a soil-borne plant pathogen, which caused devastating damaging agriculture productions worldwide. The plasticity of the pathogen has been identified with many *formae speciales* according to their host range [1]. Fusarium wilt of watermelon, caused by *F. oxysporum* f. sp. *niveum* (Fon), is one of the limiting factors for watermelon production worldwide [2,3]. During most of watermelon growing stages, Fon infects watermelon and causes annual losses of watermelon production to 10% to 15% of yield [4,5]. One of the most recommended methods for controlling this disease is breeding the resistant line; however, this is a time-consuming approach [2]. Another approach is using non-pathogenic *F. oxysporum*, which antagonises the pathogenic Fo or primes host immune response. Therefore, understanding the pathogenesis and virulence of Fo is pivotal in controlling Fusarium wilt in the field [6].

The progress of Fo introduced in their host contains four significant steps [7]. First of all, Fo requires the queue of host plants to start their invasion processes. Roots secret broad range of organic signals such as sugars and amino acids which can trigger Fo spore germination and support germ-tube growth [8]. Secondly, germ tube of Fo will then grow to root surfaces where the first contact point of pathogen-plant interaction [9]. Fo attaches the surface of root hairs and form a mycelial network and colonised the host root system [10]. Thirdly, Fo invades root cortex and vascular tissues and differentiates within xylem vessels. Fo secrets a broad range of enzymes to breakthrough the first layer of basal plant defence [11]. Finally, Fo secrets toxins and virulence factors which assist disease evolvement in the host plant [12]. All of these steps were involved with virulence factors that regulate the pathogenesis of Fo to host plants.

Degeneration of fungi is a common feature for most of fungi cultivation. The feature commonly happens through successive subculture [13]. These changes include phenotypic degeneration and virulence attenuation. Additionally, successive subcultural degeneration will affect fungal pigmentation, growth rate, morphology, rate of spore production, changes in metabolic products, and most of the time production of variant sectors [13]. Studies have demonstrated the occurrence of phenotype degeneration of *Fusarium* spp. after successive subculture [14]. In the normal state, wild-type Fo grew on media by stacking the macroconidia and forming sporodochia [15]. Reports showed there were two types of Fo colonies forming after degeneration, mycelial type (MT) and pionnotal type (PT) [16]. Mycelial type of Fo produces mycelia without pigment deposition and conidia production. In contrast, pionnotal type produces no mycelia but with a massive amount of conidia spores which forming slimy type of colonies. The development of fungi after degeneration has been described; however, the impact of degeneration on virulence remains unclear.

Although few studies have been reported the phenomenon of fungi degeneration, the major impacts of this process to virulences of *F. oxysporum* remains unclear. Therefore, we choosed *F*. *oxysporum* f. sp. *niveum* (Fon-H0103) to understand the impacts of fungi degeneration which caused by successive subculture. First of all, we replicated successive cultures for ten generations and observed the occurrences of degeneration. Secondly, we selected four different degenerated variants for the following analyse to identify impacts of degeneration to development and virulences of Fon. Lastly, we examined the cell wall-degrading enzymes activities and virulence-related genes expressions to reveal the mechanisms of fungi degeneration.

## 2. Materials and Methods

### Pathogen resources and cultivation

*Fusarium oxysporum* f. sp. *niveum* (Fon-H0103 isolate) was grown on 1/2 PDA (half-strength potato dextrose agar, potato extract 200 g L^−1^, 1% D-glucose, 2% agar). Four degenerated variants (PT2, PT3, PT11 and MT6) were derived from successive subcultures of Fon-H0103 sporodochial type (ST). Fon ST and degenerated variants were maintained with single spore culture every two weeks on Nash-PCNB plate (1.5% peptone, 2% agar, 0.1% KH_2_PO_4_, 0.5% MgSO_4_•7H_2_0, 0.1% pentachloronitrobenzene, 0.03% streptomycin and 0.1% neomycin) to avoid further degeneration [17].

### Plant materials and growth

Susceptible watermelon (Grand Baby, GB, Know-you Ltd) was used in this research. GB seeds were treated with running water at room temperature for two days. The germinated seeds were placed on water-soaked filter papers (ADVANTEC^®^, Tokyo Roshi Kaisha, Ltd., Tokyo, Japan) in the plastic petri dish for two days in the dark. The cultivated GB seedlings were then subjected to the pathogen inoculation.

### Generation of *F. oxysporum* degenerated variants

*F. oxysporum* f. sp. *niveum* (Fon ST) were cultured on half-strength PDA for one week. The fungal discs from the aged mycelia and hyphal tip, which demonstrated in the scheme (sup. figure 1), were used for successive subculture. At least ten successive subcultures were cultivated and recorded. The degenerated or sectorisation of Fon were then isolated as single spore to maintain the degenerated variants. The pattern of fungal degeneration was recorded, and the transformation rate was calculated. In the end, three pionnotal types (PT2, PT3, and PT11) and one mycelial type (MT6) were selected for further experiments.

### Phenotyping of fungus growth

Growth phenotypes of Fon ST and degenerated variants (PT2, PT3, PT11 and MT6) were examined by three types of growth indices: mycelium growth, spores production, chlamydospores formation. For mycelium growth, the single spore of Fon ST and degenerated variants was placed in the centre of half-strength PDA plate. The colony diameter was measured after seven days incubation at 28°C. To elucidate the ability of conidia spores production, a 0.5 cm^2^ fungal disc from 3-weeks growth of Fon ST and degenerated variants were cut from half-strength PDA and washed in 2 mL sterilised water to make spore suspensions. The spore suspensions were then examined under a microscope and calculated the numbers of spores by hemocytometer. Chlamydospore formation of each variant was determined according to the method described by Awuah et al. (1988) with minor modification [18]. The soil extract solution (500 mL) was prepared for inducing the chlamydospore formation by shaking 50 g of organic soil (peat moss soil BVB No.4: Bas Van Buuren Co., Ltd., Maasland, Holland) for one hour. The slurry was filtrated through a 9-cm Whatman No. 1 filter paper for clarification of the soil extract. The soil extract was sterilised by filtering through a 0.2 μm Millipore filter (Millipore Corp., Bedford, MA, USA). Five hundred μL of conidia suspension (10^4^ spores mL^−1^) was mixed with 6 mL of the sterilised soil extract and further plated on a 5-cm petri dish at room temperature for seven days. The chlamydospore formation was observed and counted by using glass hemocytometer under a microscope.

### Methods for pathogen inoculation

Fon ST and degenerated variants (PT2, PT3, PT11 and MT6) were used for inoculation tests to determine their virulence or pathogenicity. Two inoculation methods were applied. For root dipping method, susceptible watermelon GB seedlings were grown on peat moss soil BVB No.4 (Bas Van Buuren Co., Ltd., Maasland, Holland). Two weeks after growing, GB seedlings were up-rooted and cleaned with running water to reveal the seedlings’ rooting system. The second method was infested soil inoculation according to Chang et al. (2015) [2]. Fon ST and degenerated variants were cultivated in half-strength PDA, after two weeks cultivation, the Fon mycelium and spores were collected from the plate and mixed in the sterilised sand. Concentrations of Fon in infested soil were examined by plating the serially diluted soil suspensions on PCNB plates and calculate the concentrations of Fon in the stock infested soil. The stock infested soil was then diluted with peat moss soil BVB No.4 to 10^4^ spores per gram of soil. Germinated watermelon seedlings were grown in the infested soil for Fon ST and degenerated variants inoculation.

### Cell wall-degrading enzymes assay

The cell wall-degrading enzymes (CWDEs) were assayed according to the methods modified by King et al., (2009) [19]. Different variants of Fon were cultured in Bilay and Joffe’s medium with modifications (0.1% KH_2_PO_4_, 0.1% KNO_3_, 0.05% MgSO_4_·7H_2_O, 0.05% KCl, 0.02% sucrose, 0.02% glucose, 0.15% carboxymethyl cellulose, 0.15% xylan, 0.15% pectin, pH 4.0) for 7 days, the growth medium was collected to analyse the activities of cellulase, pectinase, and xylanase [20]. At the beginning of the assay, the filtered enzymes extracts from growth medium were subjected to Bradford’s protein assay to quantify the total protein concentration [21]. The CWDEs’ enzyme activities were determined by the production of reducing group after enzyme reaction with the colourimetric assay by using 3,5-dinitrosalicylic acid (DNS). The reaction solution containing carboxymethylcellulose, xylan and citric pectin were used for cellulase, xylanase and pectinase activities, respectively, with minor adjustment (pH=4.8). Activities of CWDEs were determined by analysed the production of reducing sugar after an hour reaction at 40°C.

### Real-time quantitative PCR (qRT-PCR) of virulence factors gene expression

For gene expression analysis, variants of Fon obtained from overnight PDB (potato extract 200g L^−1^, 2% D-glucose) cultures were regrown for 12 h in fresh PDB medium. Cultures of mycelium were filtered through three layers of MiraCloth (Calbiochem Corp., La Jolla, CA, USA) for nucleic acid isolation. Total RNA was extracted using the RNAzol^®^RT (Molecular Research Center, Inc., Cincinnati, OH, USA) according to the manufacturer’s procedure. Tested cDNA was prepared using the SuperScript™III RNaseH^−^ reverse transcriptase system (Invitrogen, Carlsbad, CA, USA) according to the manufacturer’s procedure. Real-time PCR was monitored on Rotor-Gene^®^ Q-Pure Detection System (Software Ver. 2, Qiagen Inc., Valencia, CA, USA) and performed using QuantiFast SYBR^®^ Green PCR Master Mix (Qiagen). Primers specific to each of the tested virulence-related genes were designed by NCBI (National Center for Biotechnology Information, USA) net program Primer-BLAST and listed in Supplement Table 1. For real-time PCR assay, a ten μl reaction mixture containing template cDNA (synthesised from 10 ng total RNA), each of 100 nM amplification primers, and 1X QuantiFast SYBR^®^ Green PCR Master Mix (Qiagen). The parameters for real-time PCR were according to the procedure of the manufacturer (polymerase activation hold at 95°C for 2 minutes, then 40 cycles of denaturing at 95°C for 15 seconds and of annealing/extension for 1 minute). After real-time PCR, melting curves (65°C to 99°C) of the PCR products were analysed to verify the specificity of the amplified fragments. Three independent experiments were used for calculating the relative expression level of each virulence-related gene.

## 3. Results

### 3.1. Successive subculture of aged mycelia induces the cultural transformation

The degeneration of Fo happens during the successive subculture. In order to understand the occurrence of degeneration and the causes, Fon culture discs from the aged mycelium and the fresh hyphal tips were used for a successive subculture (Figure 1). Results of successive culture from aged mycelium showed 10% transformation rate at first subculture and inclined toward 60% transformation rate at the 10th subculture. However, successive culture from hyphal tips showed no transformation during all subcultures.

**Figure 1.**
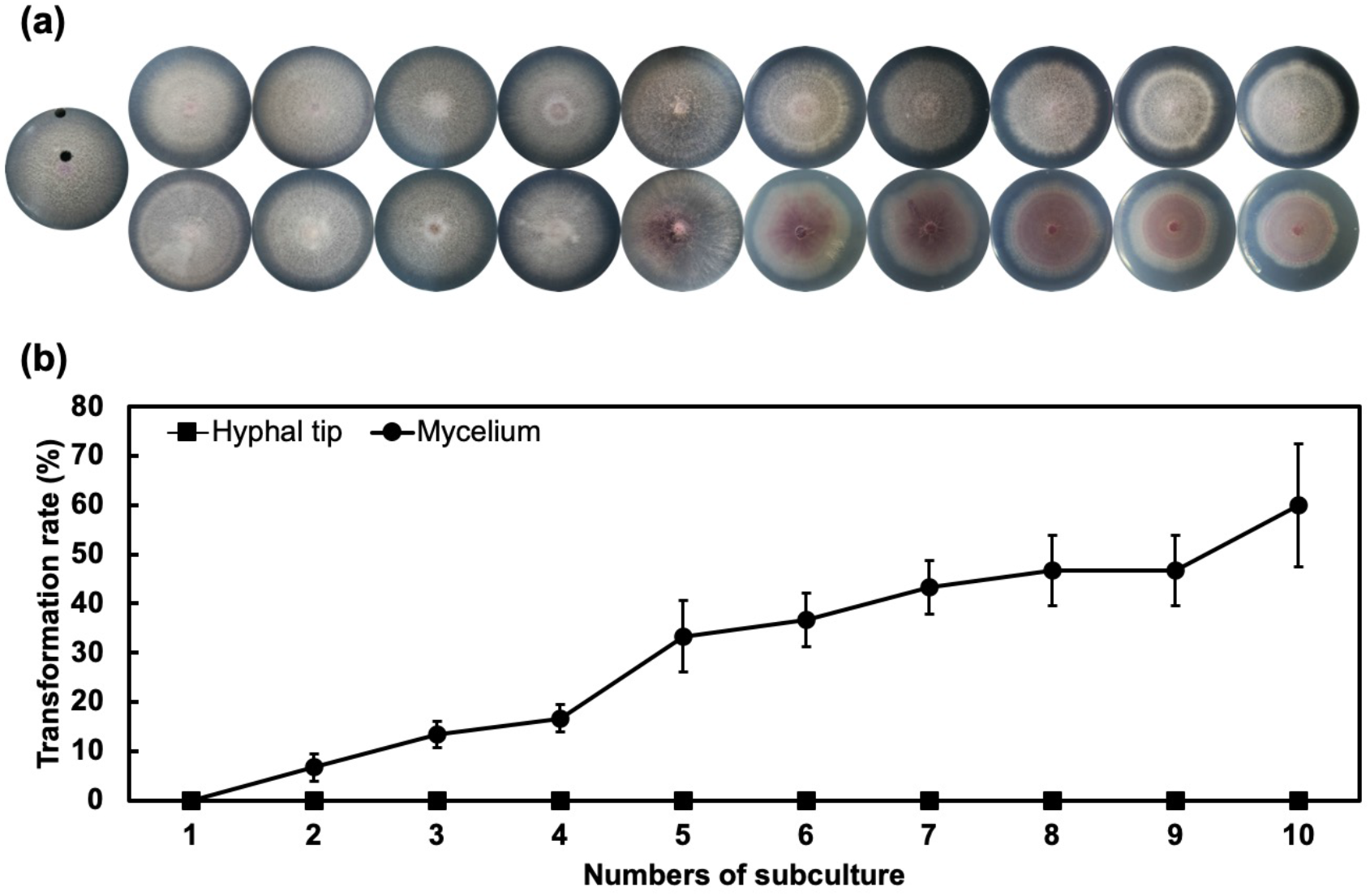
Successive subculture of aged mycelia induces transformation rates of Fon. (a) Snapshot of Fon growth at different sequential subcultures (left to right: 1st to 10th subculture) that originated from middle aged mycelium (bottom lane) and fresh hyphal tip (top lane). (b) Transformation rates of Fon from different areas of the culture plate. Error bars represent the standard error of three replicated experiments.

### 3.2. Variants of Fon demonstrated significant morphology changes

To analyse more specific changes, we then choose four degenerated candidates (include three pionnotal types PT2, PT3 and PT11, and one mycelial type MT6) for a broader range of examination to identify the effects of degeneration from the successive subculture.

Colony morphology of the Fon variants PT2 (Figure 2a), PT3 (Figure 2b), and PT11 (Figure 2c), variant MT6 (Figure 2d), and their parental Fon ST (Figure 2e) were pionnotal, mycelial, and sporodochial, respectively. When grown on PDA, the differences in growth rates between the cultures were not significant (Figure 3a). The production of conidia of all pionnotal variants was significantly higher than that of MT6 and ST-H0103 (Figure 3b), but the capacity of PT variants to produce chlamydospore was all much lower than MT6 and ST-H0103 (Figure 3c). These data indicated that the tested Fon cultures had some cultural differences in conidiation and chlamydospore production but not in hyphal growth.

**Figure 2.**
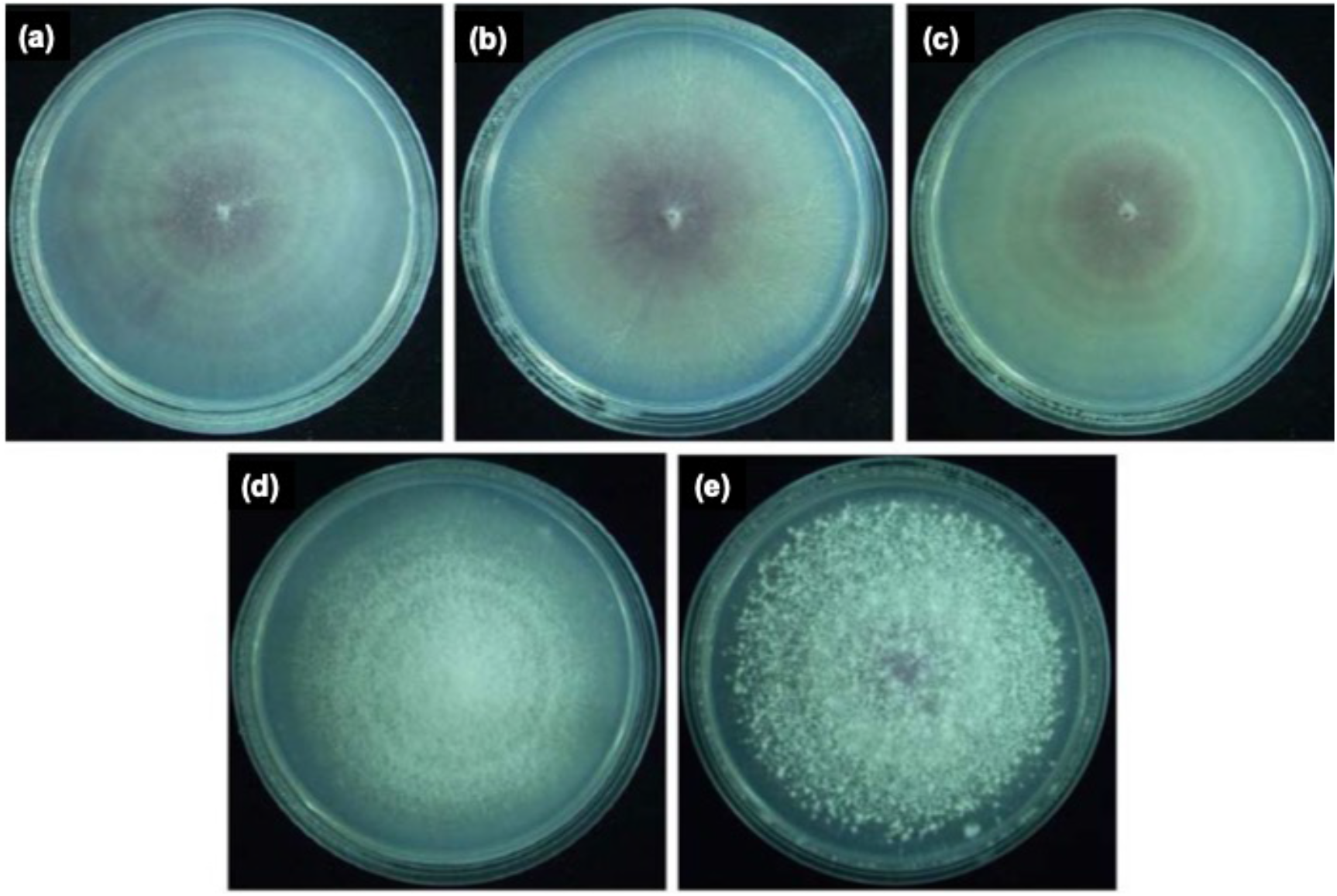
The colony morphology of the watermelon Fusarium wilt pathogen variants PT2, PT3, PT11, MT6, and their parental culture Fon ST. Variants PT2 (a), PT3 (b), and PT11 (c) belong to pionnotal type with slime surface and almost no aerial mycelia formed. Variant MT6 (d) belongs to mycelial type with abundant aerial mycelia but lack of sporodochia. Parental ST-H0103 (e) belongs to sporodochia type with abundant aerial mycelia and sporodochia.

**Figure 3.**
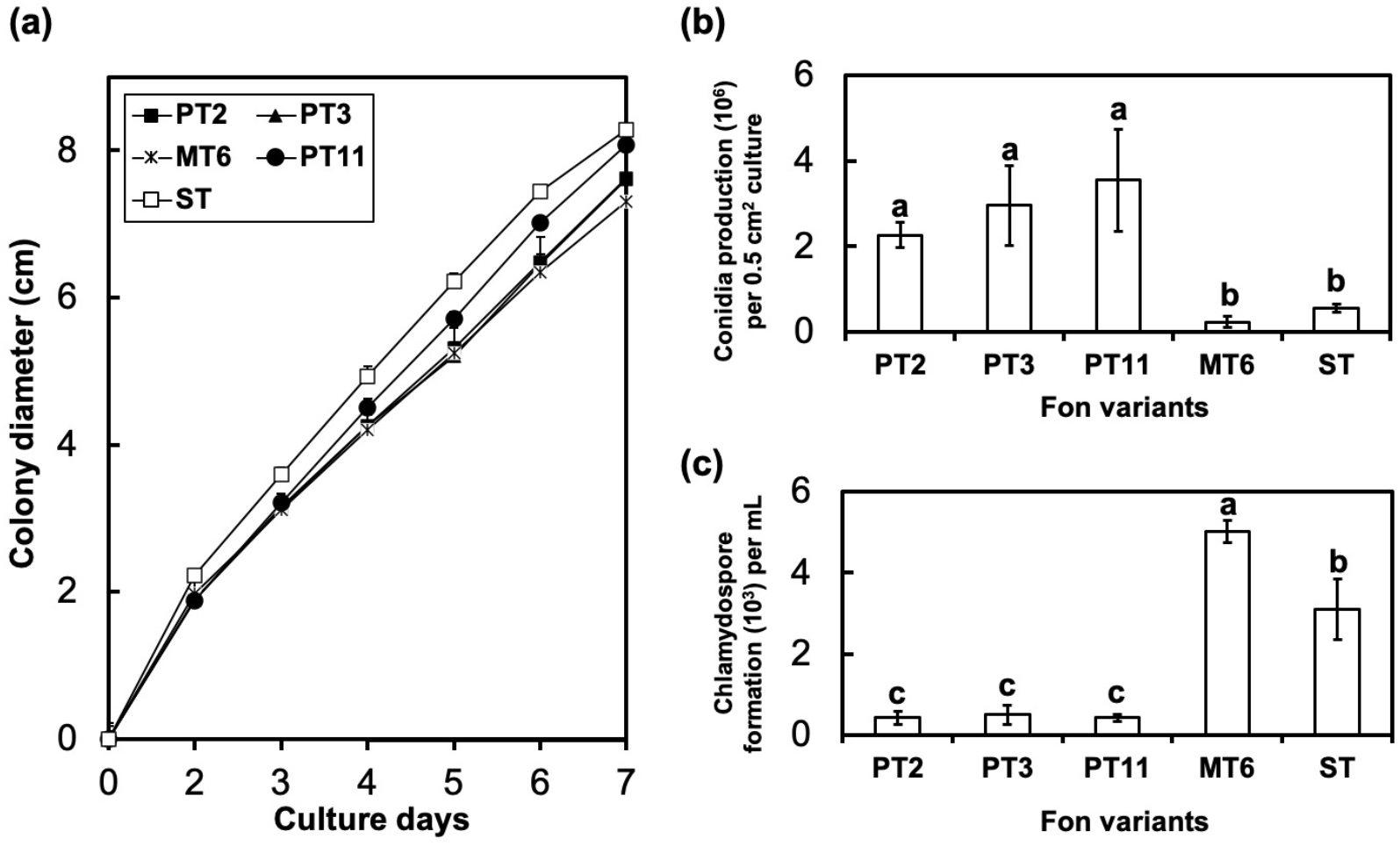
Development of colony diameter (a), conidia production (b), and chlamydospore formation (c) of the watermelon Fusarium wilt pathogen variants PT2, PT3, PT11, MT6, and their parental sporodochial culture Fon-H0103 (ST). (a) No significant statistic difference in colony diameter were observed between each culture of Fon. (b) The pionnotal variants (PT2, PT3, and PT11) produced the highest amounts of conidia, whereas the mycelial variant (MT6) produced the lowest amount of it. (c) The mycelial variant (MT6) produced the highest amount of chlamydospores, and the pionnotal variants (PT2, PT3, and PT11) produced the lowest amount of it. The values were analysed by least significant differences (LSD) test and values with different letters indicate significant difference (p < 0.05).

### 3.3. Variants of Fon reduced their virulence to susceptible watermelon

Virulences of Fon variants were tested by root-dipping and infested-soil methods. The disease progress of Fon variants were recorded with disease severity (supplement data). Virulences of variants were demonstrated by calculating the area under disease progress curve (AUDPC). The results of root-dipping showed that the ST and other variants had no significant differences in virulence (Figure 4a). However, the results of infested-soil method demonstrated that virulences of Fon variants (PT2, PT3, PT11 and MT6) were lower than Fon ST (Figure 4b). With the two inoculation system, we showed that Fon variants had lost their virulences while degeneration occurred.

**Figure 4.**
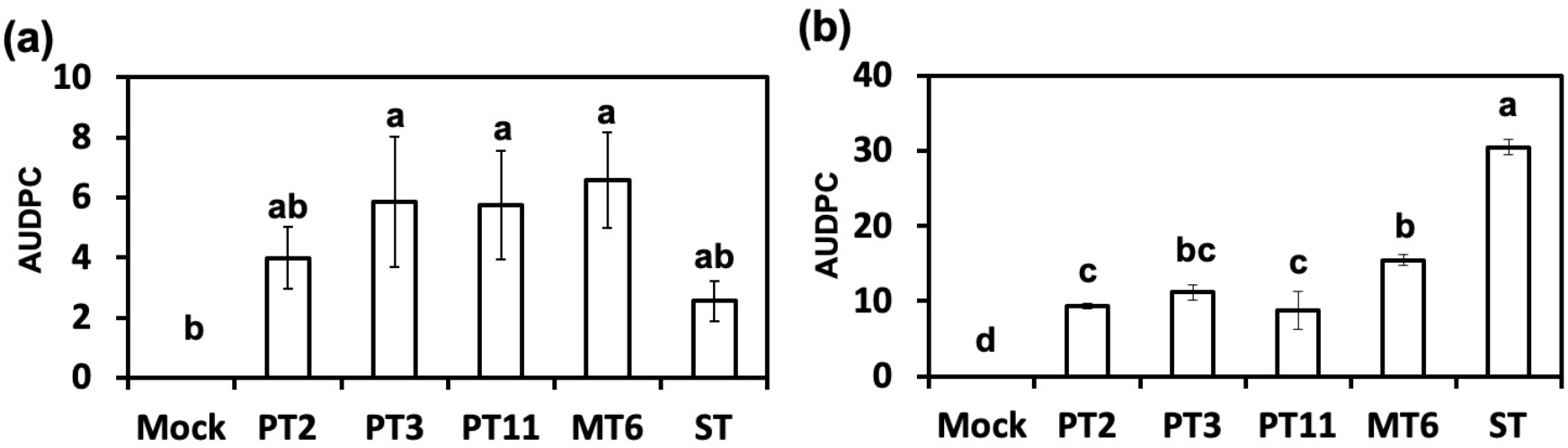
Virulence test of different Fon variants using root-dipping and infested-soil methods. The area under disease progress curve (AUDPC) of different variants through root-dipping (a) and infested-soil (b) methods was analysed. Error bars represent the standard error of three replicated experiments. Data with the same letter are not significantly different according to LSD test (p < 0.05). Mock: water inoculation control; ST: Fon sporodochial type; PT2, PT3, PT11: pionnotal type; MT6: mycelial type.

### 3.4. Activites of cell wall-degrading enzymes (CWDEs) were reduced in dedenerated variants

Following the hypothesis of how Fon variants lose their virulences, a biochemical assay for assessment of Fon variants, cell wall-degrading enzyme activity, was applied. The enzyme assays showed that the activities of CWDEs cellulase (Figure 5a), pectinase (Figure 5b), and xylanase (Figure 5c) in Fon ST were all higher than those in the variants PTs and MT6. Cellulase and xylanase activities were close to zero which were ten times lower in variants than in Fon ST. Moreover, the pectinase activities of all degenerated variants were lower than that of Fon ST. These results suggest that degeneration of Fon affects the CWDEs enzyme activity which might reduced the virulence of Fon to watermelon.

**Figure 5.**
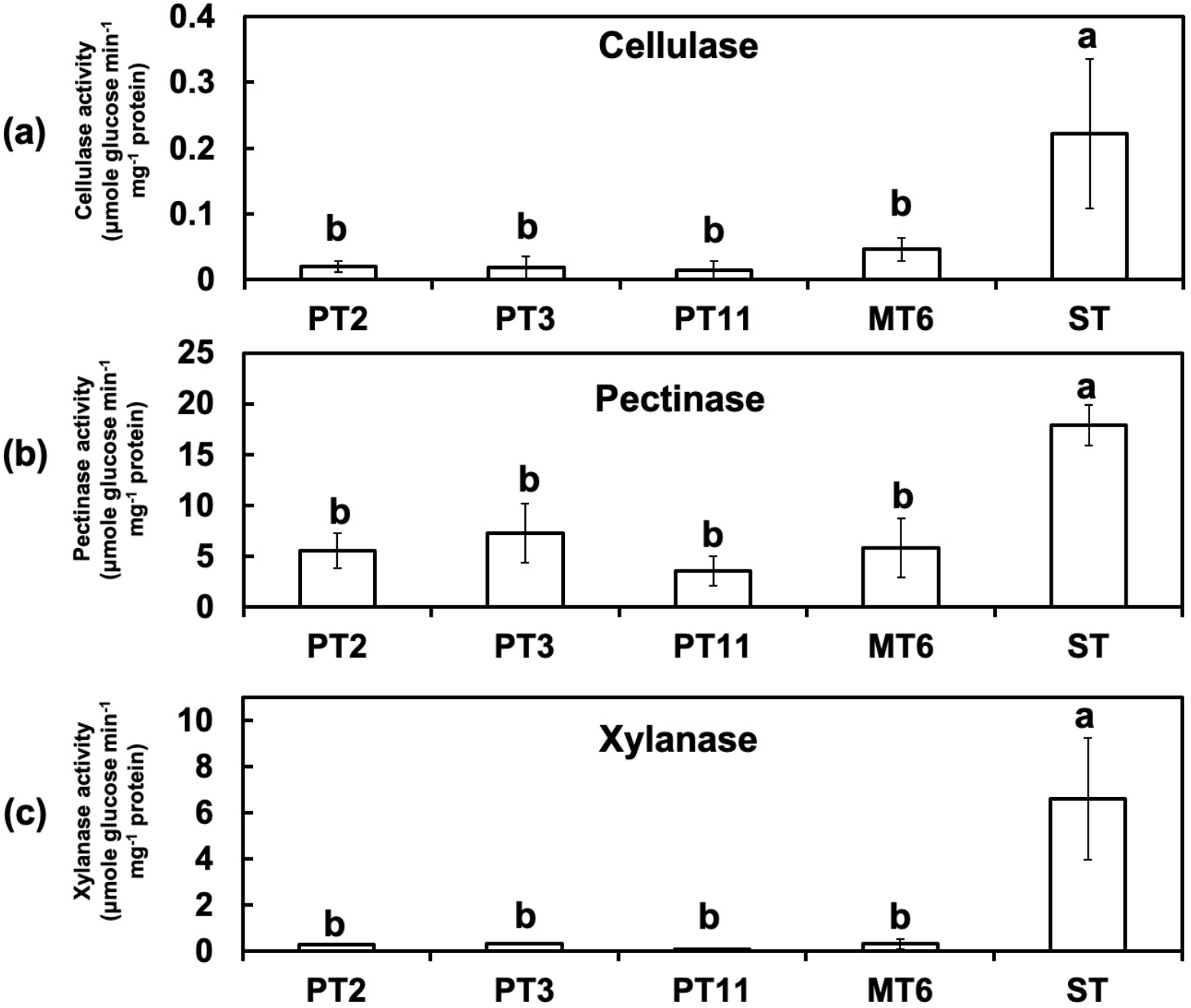
Cellulase, pectinase, and xylanase activities of different variants of Fon. Different variants of Fon culture in Bilay and Joffe’s medium with modifications were grown for 7 days, the growth medium was collected to analyse the activities of cellulase (A), pectinase (B), and xylanase (C). Bars represent the standard error. Data with the same letter are not significantly different according to LSD test (p < 0.05). ST: Fon sporodochial type; PT2, PT3, PT11: pionnotal type; MT6: mycelial type.

### 3.5 Fungus degeneration affects the expression of virulence-related genes

Expression profiles of the virulence-related genes including *fmk1* (Figure 6a), *fgb1* (Figure 6b), *pacC* (Figure 6c), *xlnR* (Figure 6d), *pl1* (Figure 6e), *rho1* (Figure 6f), *gas1* (Figure 6g), *wc1*(Figure 6h) and *fow1* (Figure 6i) in the cultures of Fon were determined by qRT-PCR analysis. Comparison with ST-H0103, relatively low level of transcripts of the virulence-related genes (*fgb1*, *pacC*, *pl1*, *rho1*, *gas1* and *wc1*) were accumulated in mycelia of all variants. On the other hand, *fmk1*, *xlnR,* and *fow1* genes expression in all variants were not consistant. Expression of *fmk1* was 2.2 times higher in PT3 than in ST, but 0.8 times lower in MT6 than in ST. PT11 showed no significant differences to ST in *xln*R and *fow*1 genes expression and was significant higher than other variants. These observations indicates that a sufficient accumulation of the virulence-related factors in mycelia seems to be required for full pathogenesis of Fon.

**Figure 6.**
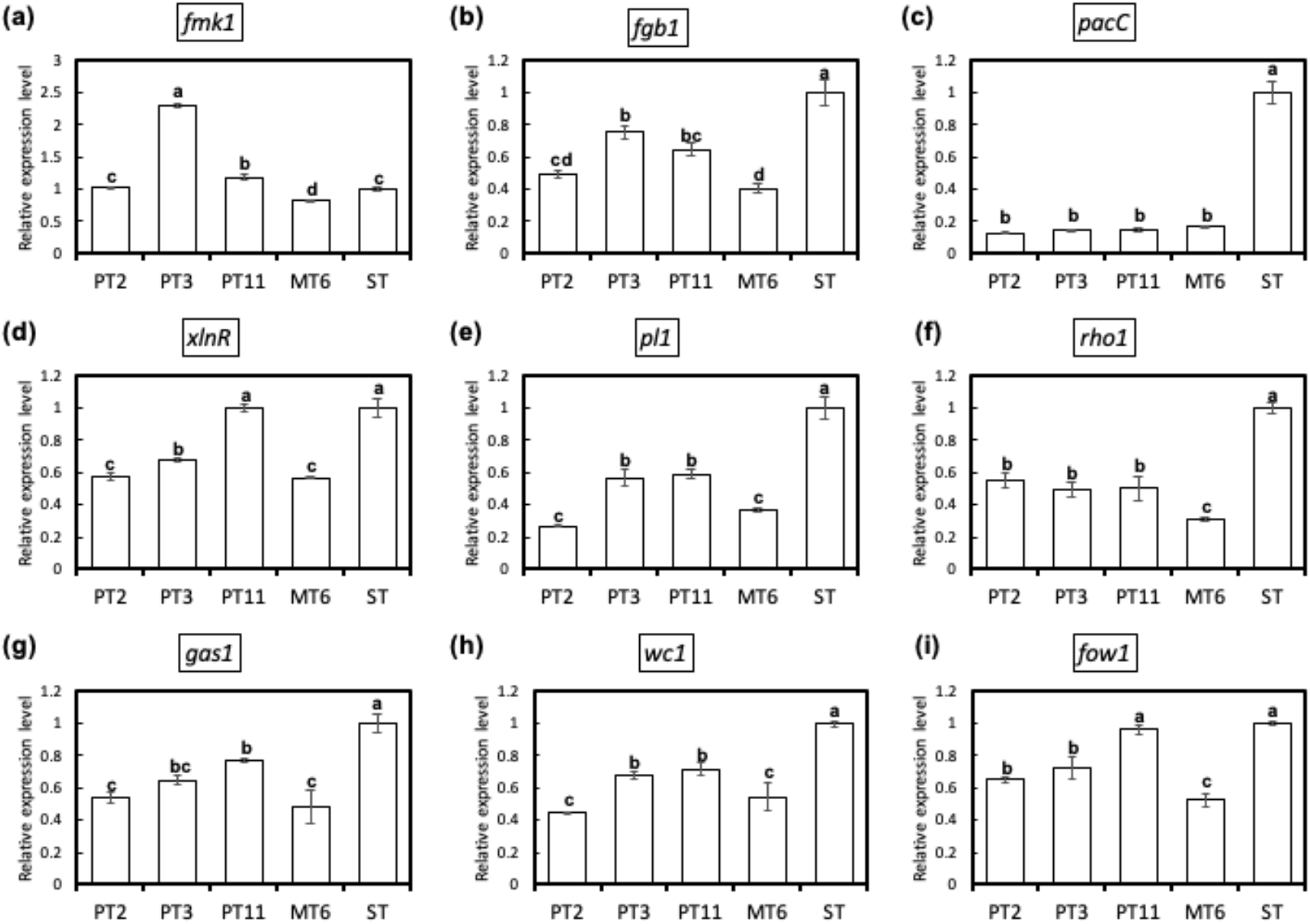
Expression analysis of virulence-related genes by real-time quantitative RT-PCR (qRT-PCR) in the watermelon Fusarium wilt pathogen Fon variants PT2, PT3, PT11, MT6, and their parental sporodochial culture ST-H0103 (ST). Quantification was achieved by normalizing the virulence-related genes *fmk1* (a), *fgb1* (b), *pacC* (c), *xlnR* (d), *pl1* (e), *rho1* (f), *gas1* (g), *wc1* (h) and *fow1* (i) to the endogenous housekeeping gene of elongation factor 1-alpha (EF1α) using the 2^−ΔΔ^ C_T_ method published by Livak & Schmittgen (2001). The formula 2^−ΔΔ^ C_T_ was used to calculate the relative transcripts of the virulence-related genes between Fon ST and variants (PT2, PT3, PT11 and MT6). The details of the virulence-related genes are listed in supplement table 1. Three independent biological replicates for each culture were used for qRT-PCR. The values were analysed by least significant differences (LSD) test and values with different letters indicate significant difference (p < 0.05).

## 4. Discussion

The degeneration of fungi is a common phenomenon in many fungus cultivation. Fo occurs morphological mutations, especially when these pathogens are cultured on carbohydrates-rich media, and the impairment in pathogenicity is frequently accompanied [15]. Degeneration in the subculture of *F. compactum* and *F. acuminatum* occurred more frequent when using single macroconidia spore for ten generations than using hyphal tips [14]. Interestingly, we have demonstrated that successive subculture of aged mycelia has higher transformation rate than of hyphal tips (Figure 1). The results suggest that the occurrence of degeneration is varied from different stages of Fo.

The changes are usually reversible while fungus was grown back to the natural habitat environment instead of artificial [22]. However, morphological changes of the variants, PT2, PT3, PT11, and MT6, which transformed from their parental isolate ST are irreversible (data not shown). Additionally, these variants demonstrated significant morphology differences compared with ST (Figure 2). The irreversible changes make these potential variants candidates for elucidating effects of cultural degeneration.

Our research demonstrated that the growth rates between Fon ST and degenerated variants on the PDA medium were similar. The observation in the growth rate of the morphological variants of Fon is consistent with the finding in *F. oxysporum* f. sp. *apii* [18] but not with that in *F. graminearum* [23]. Additionally, the production of conidia spore and formation of chlamydospore showed significant differences between Fon-H0103 ST and degenerated variants. The mass production of conidia spore in PT variants and induction of MT variant chlamydospore formation suggests fungus degeneration may affect the pathogenicity of Fon.

Therefore, we assessed the virulence of Fon ST and degenerated variants. Our results showed that the degenerated variants delayed or attenuated the disease progress in watermelon with the infested-soil assay. However, the watermelon Fusarium wilt development of degenerated variants showed no significant differences with Fon ST through root-dipping method. The root-dipping method increases the opportunity for Fon to contact roots surfaces and the root system often was wounded through up-rooted methods [24]. In other words, the root dipping method has eliminated the first layer of watermelon defence system. Hence, we speculated that the degenerated variants might cause loss of virulence factors which were related to the stage of host-recognition and colonisation.

Plant cell wall, the front layer for the plant defence system, composes polysaccharide-rich elements such as cellulose, hemicellulose, and pectin [25]. The cross-linking of polysaccharides and other intermediates such as phenolic compounds make the cell wall rigged, which prevent the intruding attempt from pathogens [26]. In watermelon, the reinforcement of the cell wall after pathogen inoculation is thought to be an essential character for breeding the Fusarium-resistant lines [2]. CWDEs are practical tools for pathogenic microorganisms to invade host plants [12]. The induction of pectinase and xylanase of *F. oxysporum* f. sp. *ciceris* are positively correlated to the disease development in chickpea [27]. However, ameliorating an individual CWDE did not show a significant effect on virulence which may because of functional redundancy [3]. Knockout of *XlnR*, a transcription activator for xylanase genes, did not affect the virulence of *F. oxysporum* f. sp. *lycopersici* in tomato [28]. In the present study, the activities of cellulase, pectinase and xylanase were decreased in degenerated variants compared with Fon ST. The phenomenon of losing all arsenals for breaking plant cell wall might reduce the virulence of Fon to watermelon.

Forward and reverse genetics approaches have been used to identify virulence-related factors for pathogenesis. Most of the virulent factors were implicated in fungal cell wall biosynthesis, signal transduction processes. To understand the possible mechanism on the pathogenesis of Fon, we performed qRT-PCR to investigate the correlation between the virulence-related factors and virulence. Gas1, a β-1,3-glucanosyltransferases, a member of glycosylphosphatidylinositol-anchored glycoproteins, has been implicated as a virulence-related factor on the pathogenesis of *F. oxysporum* f. sp. *lycopersici* [29]. In addition, Gas1 is required for cell wall biosynthesis and morphogenesis of *F. oxysporum* f. sp. *lycopersici*, and the pathogen mutant deleting functional *gas1* exhibited the structural alterations in the cell wall. The present study showed that the gene expression levels of *gas1* in four variants were all significantly lower than that in their parental culture ST. The data support that *gas1* may be the most functionally important among the tested virulence-related factors in Fon pathogenesis. The virulence-related factors, G-protein β subunit Fgb1 [30], mitochondrial protein Fow1 [31], and pH response transcription factor PacC [32], have been proved to be essential for controlling the virulence of *F. oxysporum*. The effects of individuals of virulence-related factors on pathogenicity in the Fon cultures seem to be complementary to each other. In addition, less activity of CWDEs was detected in the four variants, supporting that the morphological variants in overall do not have enough weapons to attack the watermelon host. The data indicate that sufficient accumulation of virulence-related factors and CWDEs is very important for fully expressing the pathogenicity of Fon.

The genetic variation frequently observed in *F. oxysporum* has been attributed to active transposable elements [33]. Although the cultures tested in this study showed distinct variations, e.g. cultural and pathogenic variability, the genetic diversity between these cultures were not very high because the DNA fingerprinting patterns between the morphologically different cultures by RAPD (random amplification of polymorphic DNA) assays with 48 different random primers (Supplement Figure 1) were almost identical. Our results of virulence-related genes expression and CWDEs enzyme activities demonstrated significant differences between the variants and their parental ST. These results sugges that expression levels of the tested virulence-related genes and active CWDEs in the variants were insufficient to support their ability to maintain full virulence. Furthermore, the transcript levels of the virulence-genes (*fgb1*, *pacC*, *pl1*, *rho1, gas1* and *wc1*) in the mycelia of the morphological variants, were all significantly lower than in those of their parental culture ST. The data seem to be able to explain why the virulence of the variants was significantly lower than Fon ST by infested-soil method. In addition, it is worth noting that the tested virulence-genes and CWDEs may play a role in Fon pathogenesis on watermelon but they seem to be functional complementary.

It is still far from understanding of the underlying roles of individual virulence-factors during pathogenesis. Further studies devoted to whole transcriptome profiling of Fon during its pathogenesis may speed up the understandings of the function and mechanism of virulence-factors involving pathogenicity individually. Additionally, the whole transcriptome profile of Fon shall provide clues to uncovering mechanisms of loss-of-virulence during cultural degeneration.

## Supplementary Materials

**Supplement Figure 1.**
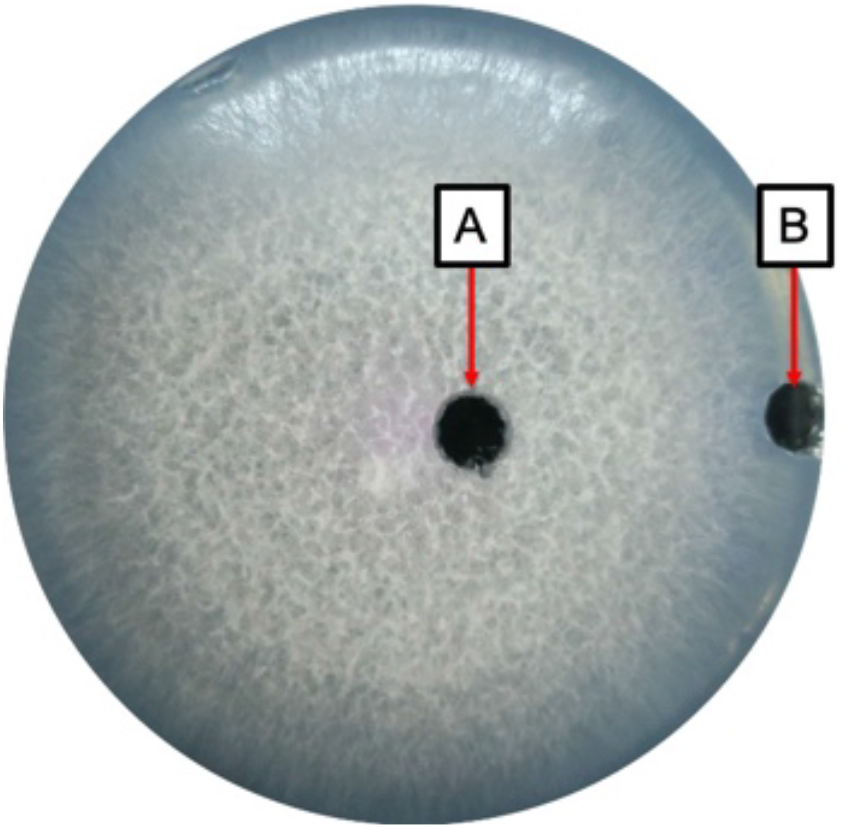
Scheme of Fon successive cultural sources which are aged mycelium (A) and hyphal tip (B).

**Supplement Figure 2.**
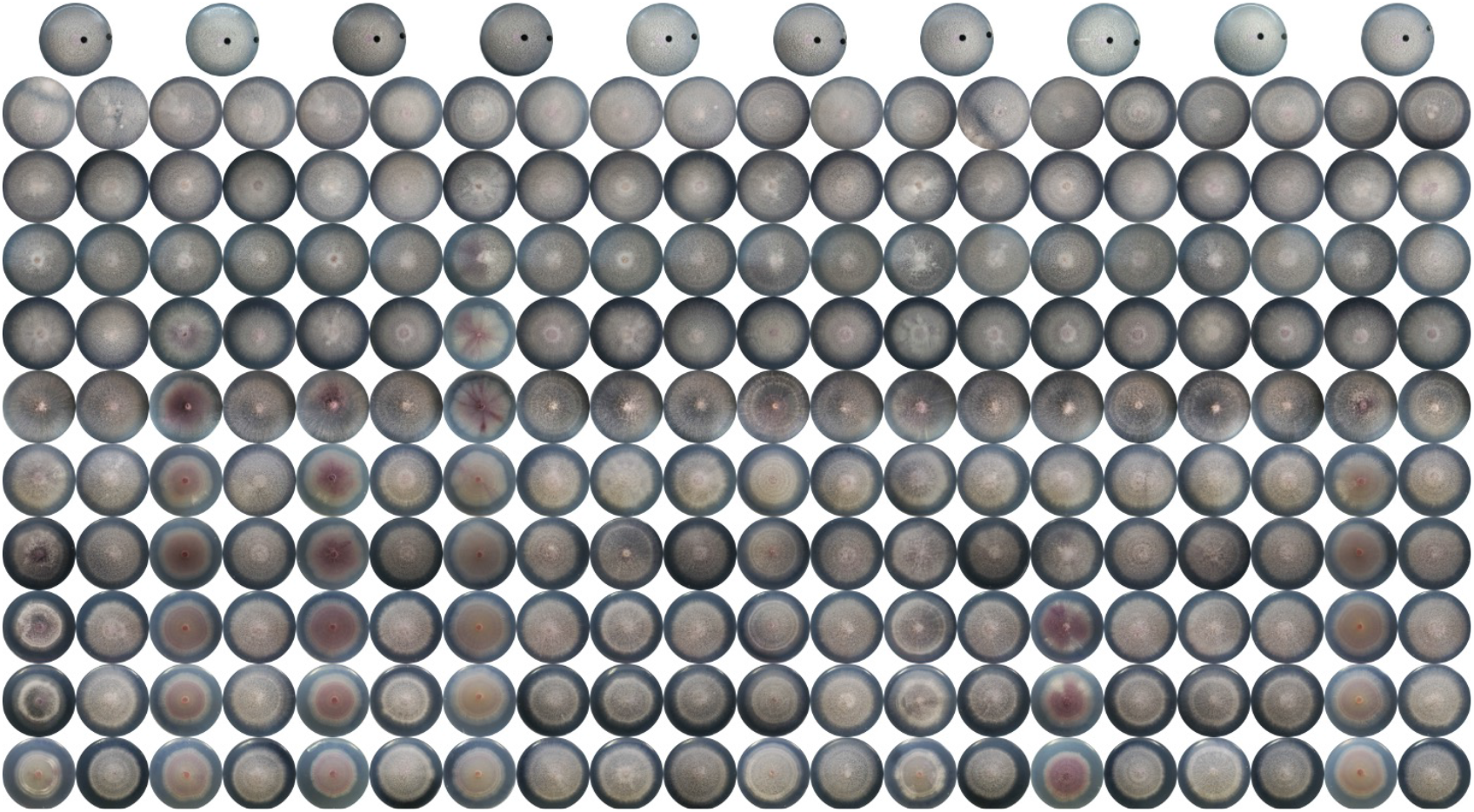
Snapshots of Fon successive culture phenotype. Fon were grown on half-strength PDA and successive cultured for 10 generations. Each successive subculture was aligned in two lanes. Every two lanes indicated cultures from one origin where cultural discs were from aged mycelium (left lane) and hyphal tip (right lane).

**Supplement Figure 3.**
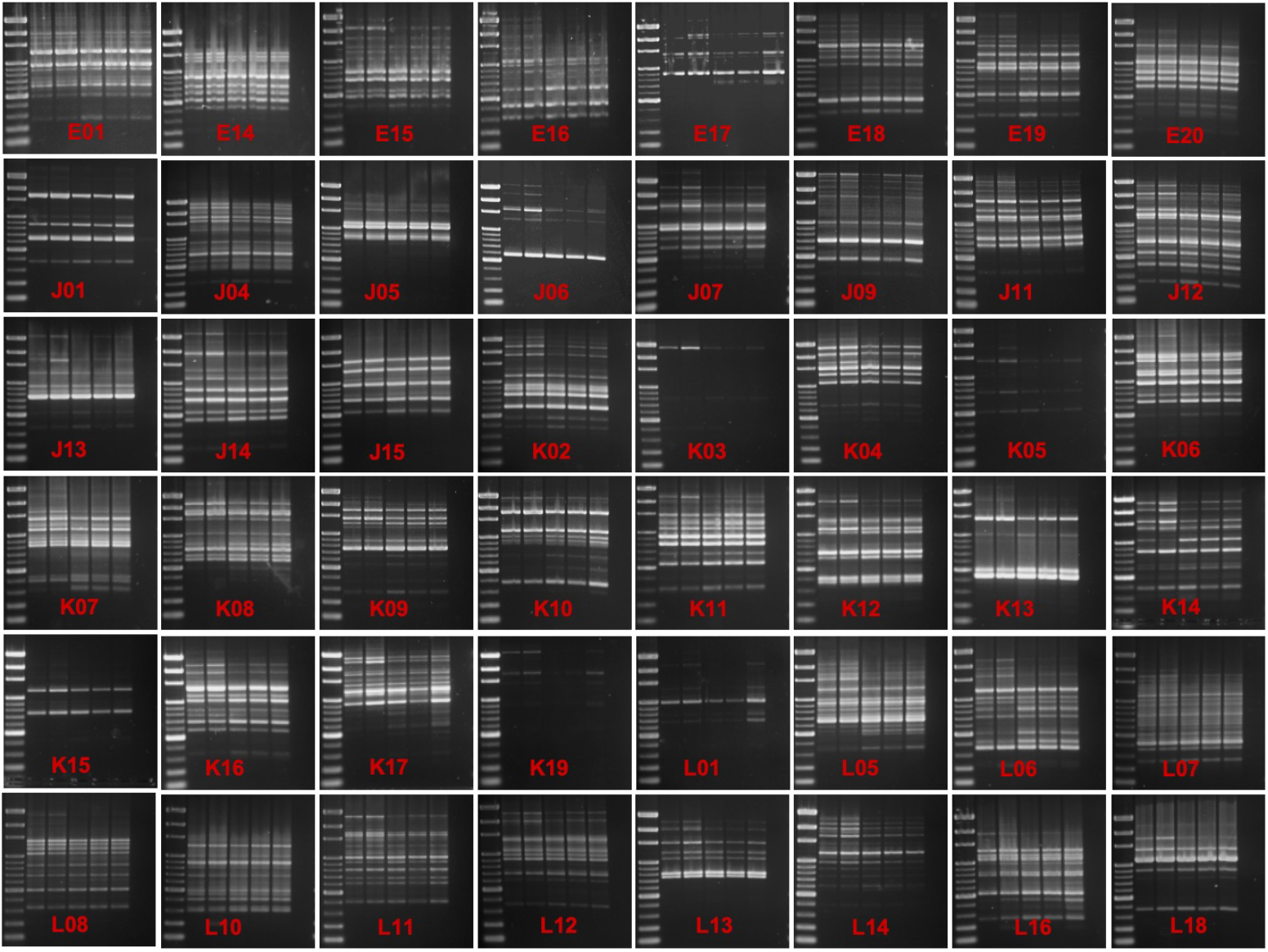
The random amplified polymorphic DNA (RAPD) patterns of different *F. oxysporum* f. sp. *niveum* (Fon-H0103) variants. Lanes of each gel figures from left to right are DNA molecular marker (100 bp marker, IDBio Ltd, Taiwan), Fon variants PT2, PT3, PT11, and MT6, and Fon ST, respectively. Total 48 random primers were used and indicated in figures.

**Supplement Table 1.**
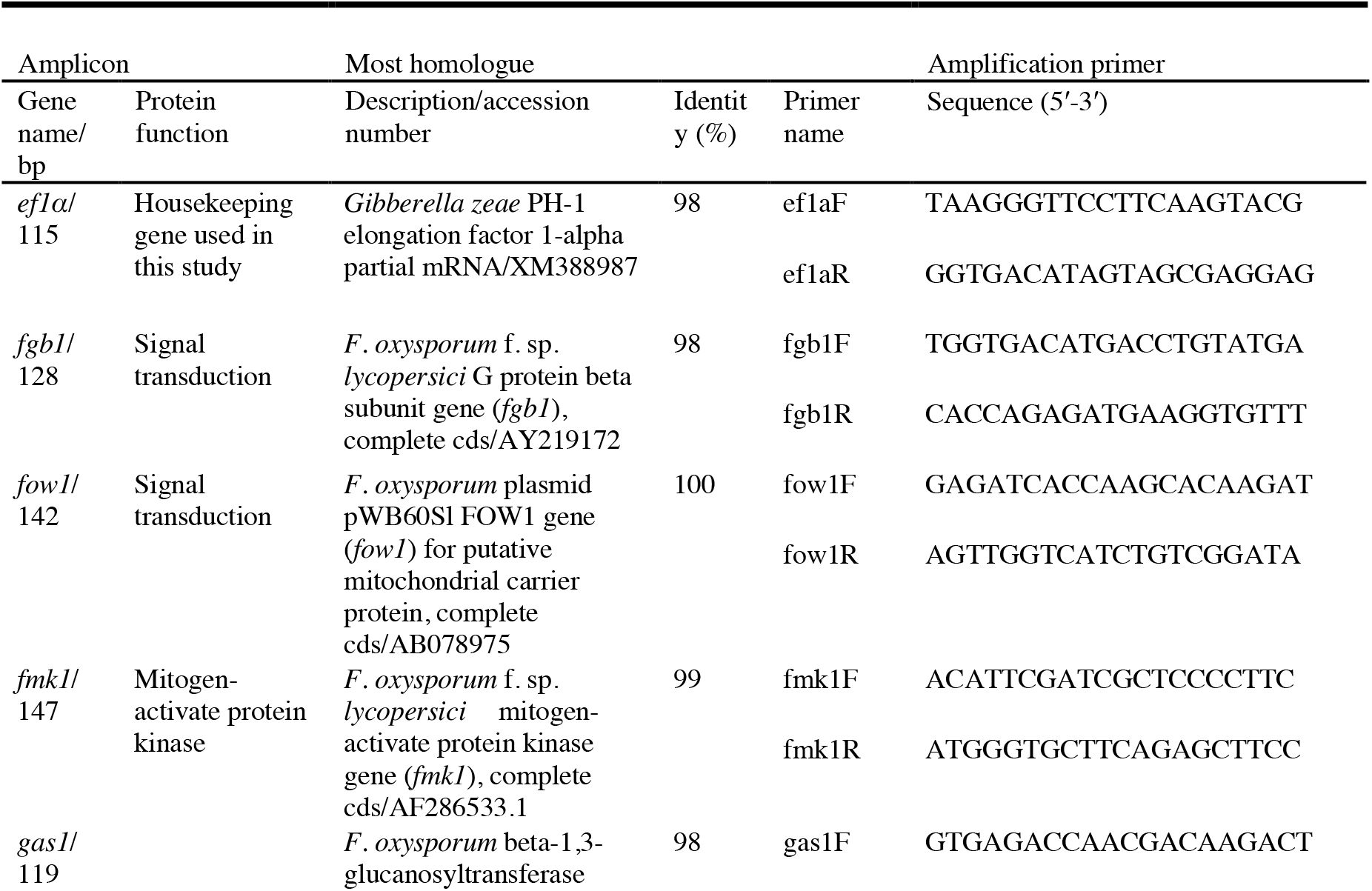
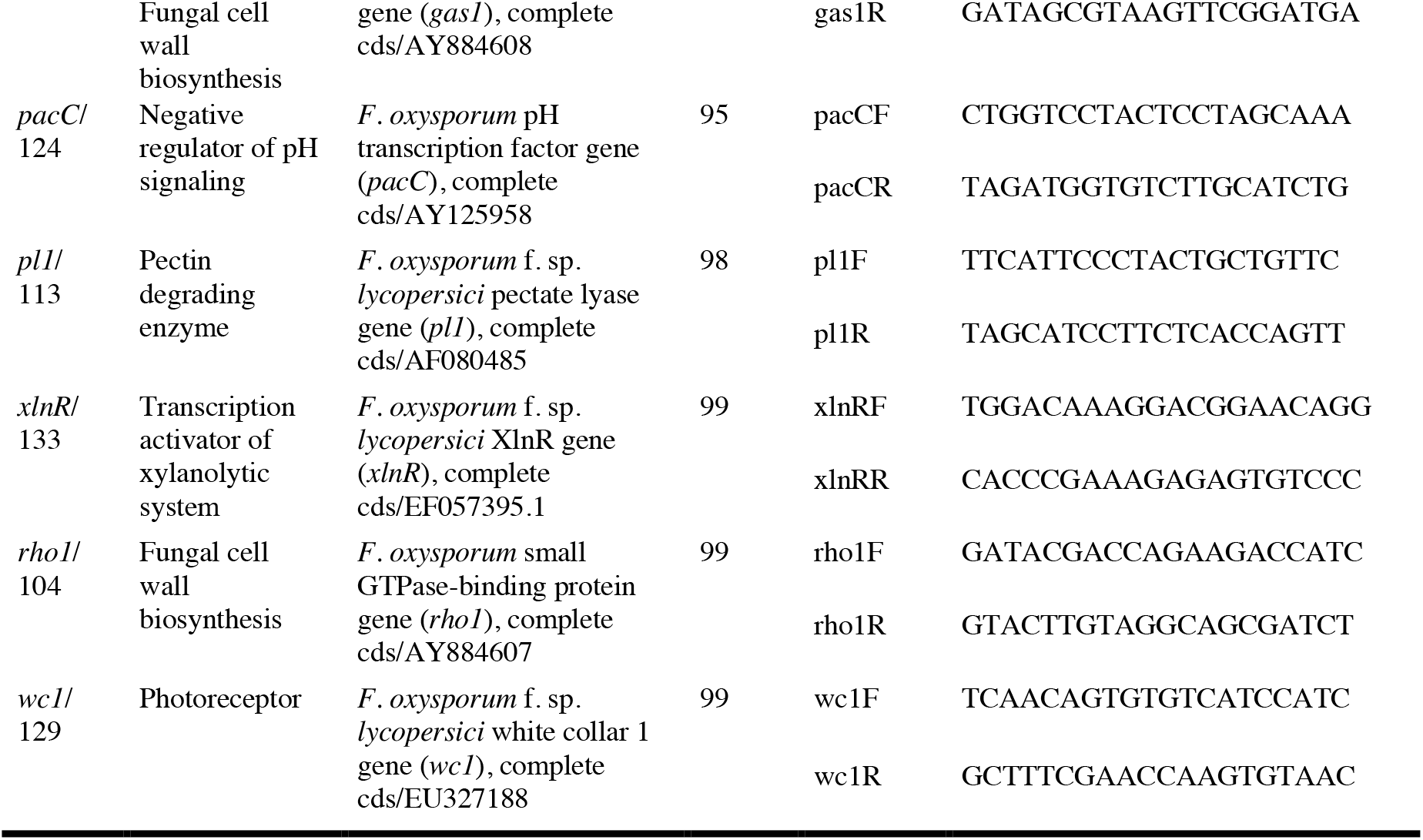
Designed primers for virulence genes expression

## Author Contributions

Conceptualization, T.-H. Chang, Y.-H Lin and Y.-L. Wan; methodology, T.-H. Chang, Y.-L. Wan and Y.-H. Lin; Software, T.-H. Chang and Y.-H. Lin; validation, Y.-L. Wan, Y.-H. Lin and T.-H. Chang; formal analysis, T.-H. Chang and Y.-L. Wan; investigation, Y.-L. Wan and T.-H. Chang; resources, K.-S. Chen, J.-W. Huang and P.-F. L. Chang; data curation, T.-H. Chang and Y.-H. Lin; writing—original draft preparation, Y.-H. Lin and T.-H. Chang; writing—review and editing, T.-H. Chang and P.-F. L. Chang; visualization, T.-H. Chang; supervision, P.-F. L. Chang and J.-W. Huang; project administration, P.-F. L. Chang and J.-W. Huang; funding acquisition, P.-F. L. Chang, J.-W. Huang and K.-S. Chen. All authors have read and agreed to the published version of the manuscript.

## Funding

This research was funded by Bureau of Animal and Plant Health Inspection and Quarantine, Council of Agriculture (Taiwan) under grant numbers under grant numbers 93AS-1.9.2-BQ-B1(1), 94AS-13.3.2-BQ-B1(6), 96AS-4.1.2-IC-I1(2), 97AS-4.1.2-IC-I1(6) and 98AS-4.1.1-IC-I1(1); by National Science Council (Taiwan) under grant numbers 98-2313-B-005-025-MY3, 99-2622-B-005-006-CC2 and 101-2313-B-005-028-MY3; by the Ministry of Education (Taiwan) under the ATU plan; and also by National Chung Hsing University (Taiwan, R.O.C.). This work is also supported in part by the “Innovation and Development Center of Sustainable Agriculture” from The Featured Areas Research Center Program within the framework of Higher Education Sprout Project by the Ministry of Education (MOE) in Taiwan.

## Conflicts of Interest

The authors declare no conflict of interest.

## Notes

### Competing Interest Statement

The authors have declared no competing interest.

